# Cardiac structure and function in patients with schizophrenia taking antipsychotic drugs: an MRI study

**DOI:** 10.1101/589093

**Authors:** Toby Pillinger, Emanuele F. Osimo, Antonio de Marvao, Alaine Berry, Thomas Whitehurst, Ben Statton, Marina Quinlan, Stefan Brugger, Ali Vazir, Stuart A. Cook, Declan P. O’Regan, Oliver D. Howes

## Abstract

Cardiovascular disease (CVD) is a major cause of excess mortality in schizophrenia. Preclinical evidence shows antipsychotics can cause myocardial fibrosis and myocardial inflammation in murine models, but it is not known if this is the case in patients. We therefore set out to determine if there is evidence of cardiac fibrosis and/or inflammation using cardiac MRI in medicated patients with schizophrenia compared with matched healthy controls. 31 participants (14 patients and 17 controls) underwent cardiac MRI assessing myocardial markers of fibrosis/inflammation, indexed by native myocardial T1 time, and cardiac structure (left ventricular (LV) mass) and function (left/right ventricular end-diastolic and end-systolic volumes, stroke volumes, and ejection fractions). Participants were physically fit, and matched for age, gender, smoking, blood pressure, BMI, HbA1c, ethnicity, and physical activity. Compared with controls, native myocardial T1 was significantly longer in patients with schizophrenia (effect size, d=0.89; p=0.02). Patients had significantly lower LV mass, and lower left/right ventricular end-diastolic and stroke volumes (effect sizes, d=0.86-1.08; all p-values <0.05). There were no significant differences in left/right end-systolic volumes and ejection fractions between groups (p>0.05). These results suggest an early diffuse fibro-inflammatory myocardial process in patients that is independent of established CVD-risk factors and could contribute to the excess cardiovascular mortality associated with schizophrenia. Future studies are required to determine if this is due to antipsychotic treatment or is intrinsic to schizophrenia.

## Introduction

Compared with the general population, people with schizophrenia are both twice as likely to have a diagnosis of cardiovascular disease (CVD), and die as a consequence of CVD.^1^ The mortality gap between individuals with schizophrenia and the general population is growing,^2^ suggesting the need for improved understanding of the factors underlying cardiovascular disease in this group.

In various cardiac diseases, myocardial fibrosis predicts both cardiovascular and all-cause mortality.^3^ Moreover, myocardial fibrosis is implicated in arrhythmogenic sudden cardiac death,^3^ a cause of mortality up to 5-times more likely in patients taking antipsychotic drugs compared with the general population.^4^ Preclinical evidence shows that dopamine D2/3 receptor activation plays a key role in regulating cardiac function^5^ and that antipsychotics impair this by blocking dopamine D2/3 receptors.^6^ Specifically, antipsychotics have been found to reduce cardiac myocyte viability,^7^ increase myocyte apoptosis^6^ and induce autophagy of cardiac myocytes,^8^ potentially leading to cardiac fibrosis.^9^ Antipsychotics are also associated with development of inflammatory lesions and lymphocytic infiltrates within the myocardium.^10, 11^ D3 knock-out mice show increased interstitial fibrosis in cardiac myocyte samples,^12^ and antipsychotics have also been found to induce interstitial and endocardial fibrosis, as well as myolysis and other changes seen in dilated cardiomyopathy.^7,9^ Moreover, antipsychotic treatment is associated with impaired cardiac function in preclinical models, including reduced left ventricular ejection fraction.^9^ Antipsychotics have other potentially cardio-toxic effects via potassium (hERG) channel blockade^13^ and the induction of myocardial oxidative stress and inflammation,^10^ processes which are associated with the development of myocardial fibrosis and functional myocardial impairment.^14^

Despite the wealth of evidence that individuals with schizophrenia are at increased risk of cardiovascular disease, there has been relatively little research directly examining cardiac function in schizophrenia. The studies that do exist have used transthoracic echocardiography (TTE) to assess cardiac function.^15–18^ These have typically shown reductions in left ventricular (LV) ejection fraction in schizophrenia with large effect sizes (d = 0.92-1.17), even in the absence of cardiac symptoms or diagnosed CVD. However, to our knowledge there have been no prior studies examining for myocardial tissue alterations characteristic of a fibrotic/inflammatory process in patients taking antipsychotic drugs, or of cardiac function using MRI, the gold standard for assessing cardiac function.^19–22^ In view of this, using cardiac MRI we set out to test the hypothesis that, compared with matched healthy controls, individuals with schizophrenia receiving antipsychotic treatment would show evidence of tissue alterations characteristic of myocardial fibrosis and inflammation, indexed by native myocardial T1 time. As secondary analyses, we also set out to examine if patients also showed changes in cardiac structure (LV mass) and function (e.g. ventricular end-diastolic and stroke volumes).

## Methods

### Subjects

A total of 17 patients with schizophrenia (SCZ) were recruited from community mental health services from the South London and Maudsley NHS Foundation Trust, London, UK. A total of 17 healthy volunteers (HV) were also recruited. We recruited HV matched 1:1 to SCZ for age (+/− 3 years), ethnicity (1:1 matching of people of Caucasian and non-Caucasian ethnicity), gender (1:1 matching of male and female), and BMI (+/− 2 kg/m^2^ of each other). Inclusion criteria for SCZ were: an ICD-10/DSM-IV diagnosis of schizophrenia, and no history or family history of other psychiatric disorders. Exclusion criteria for all participants were: age <18 or >65 years, pregnancy or breastfeeding, a past medical history of, or current/past treatment for, any physical health condition that might impact cardiac function (e.g. diabetes mellitus, dyslipidaemia, and hypertension (see eAppendix 1)), and history of past/current substance abuse, including alcohol. Exclusion criteria for HV included a personal or first-degree family history of a psychiatric disorder. Subjects were excluded if they had contraindications to MR imaging. Written informed consent was obtained from all volunteers and the study was approved by a local research ethics committee.

### Assessment of participants

Physical assessment, blood tests, study questionnaires, and cardiac MRI were all performed during the same study visit. A diagnosis of schizophrenia and exclusion of other psychiatric co-morbidity was confirmed using the Structured Clinical Interview for DSM-IV.^23^ All measurements were performed by physicians at the study centre (Hammersmith Hospital, London, UK). Brachial blood pressure measurements were performed following 5 minutes’ rest in accordance with European Society of Hypertension guidelines (eAppendix 1) using a validated oscillometric device (Omron M7, Omron Corporation, Kyoto, Japan). The first of 3 measures was discarded and the second 2 values averaged. Physical activity grading was based on the Copenhagen City Heart Study Leisure Time Physical Activity Questionnaire (eAppendix 1). Categories of activity were based on participants’ level of activity over the preceding 12 months, ranging from level 1 (almost entirely sedentary) to level 4 (>5 hours of exercise per week). Plasma HbA1c was analysed using the Tosoh G8 HPLC Analyzer (Tosoh Bioscience, San Francisco). Chlorpromazine equivalent doses and dose-years were calculated as described by Andreasen and colleagues (eAppendix 1).

### Cardiac MRI Protocol

For all participants, CMR imaging was performed on a 3T Magnetom Prisma (Siemens Healthcare, Erlangen, Germany) using a combination of an 18-element body matrix coil and 12 elements of the 32-element spine coil. Myocardial tissue alterations were assessed in line with consensus guidelines (eTable 1 and eAppendix 2) by measuring the longitudinal relaxation time constant of the myocardium (native myocardial T1 time) using a Modified Look-Locker Inversion recovery (MOLLI) sequence, 5s(3s)3s variant in a mid-ventricular short axis slice during breath-hold in end-expiration. The following imaging parameters were used for this electrocardiogram gated single-shot balance steady state free precession sequence: slice thickness 8mm; flip angle 35°; echo time 1.12ms; repetition time 280.56ms; matrix 144×256; pixel size 1.4 × 1.4mm; minimum T1 100ms; T1 increment 80ms; acquisition window 167ms; GRAPPA 2 parallel imaging acceleration factor. Circle Cardiovascular Imaging (CVI), Calgary, Canada, version 5.6.3 T1 mapping software was utilised with a Siemens recommended correction factor of 1.035 applied. Each slice was divided into 6 segments as per the American Heart Association model (figure 1 and eAppendix 2). An epicardial and endocardial erosion offset of 10% was applied to the contours to ensure only myocardium was included. After visual inspection of all segments and exclusion of those segments with evidence of artefact (e.g. owing to unintended motion effects), a mean T1 relaxation time for the remaining segments of each slice was calculated.

**Table 1:** Subject clinico-demographic characteristics (n = 31). Continuous data are presented as mean ±standard deviation where normally distributed, and as median (interquartile range) where not. There were no significant group differences. MWU: Mann Whitney U test.

|  | Patients | Healthy Volunteers | Statistic |
| --- | --- | --- | --- |
| **Age (years)**<br>Mean ±SD | 39.93 ±13.08 | 37.88 ±12.19 | t = 0.45; df = 29<br>p = 0.66 |
| **Mean Arterial Pressure (mm Hg)**<br>Median (IQR) | 93.89 (7.42) | 89.43 (10.56) | MWU<br>p = 0.47 |
| **Systolic Blood Pressure (mm Hg)**<br>Mean ±SD | 125.23 ±13.15 | 123.00 ±13.43 | t = 0.46; df = 29<br>p = 0.65 |
| **Diastolic Blood Pressure (mm Hg)**<br>Mean ±SD | 77.82 ±8.81 | 79.08 ±10.48 | t = -0.36; df = 29<br>p = 0.73 |
| **HbA1c (mmol/mol)**<br>Median (IQR) | 36.00 (12.25) | 37.00 (5.50) | MWU<br>p = 0.65 |
| **Body Mass Index (kg/m<sup>2</sup>)**<br>Mean ±SD | 25.06 ±4.37 | 26.82 ±5.07 | t = 1.03; df = 29<br>p = 0.31 |
| **Physical Activity Score**<br>Mean ±SD | 2.41 ±1.00 | 2.01 ±0.86 | t = 0.96; df = 29<br>p = 0.35 |
| **Body Surface Area (m<sup>2</sup>)**<br>Mean ±SD | 1.97 ±0.41 | 2.00 ±0.21 | t = 1.96; df = 29<br>p = 0.76 |
| **Male/Female** | 7/7 | 7/12 | Fisher's Exact<br>p = 0.50 |
| **Smoker/non-smoker** | 8/6 | 6/11 | Fisher's Exact<br>p = 0.29 |
| **Caucasian/non-caucasian** | 7/7 | 10/7 | Fisher's Exact<br>p = 0.73 |
| **Chlorpromazine Equivalent Dose (mg)**<br>Median (IQR) | 398.64 (364.00) | n/a | n/a |
| **Chlorpromazine Dose Years**<br>Median (IQR) | 1.13 (4.32) | n/a | n/a |

A standard clinical protocol for assessing biventricular function and volumes was followed according to published international guidelines (eTable 1 and eAppendix 2) and blind to diagnostic group. Volumetric analysis of the cine images was performed using CMRtools (Cardiovascular Imaging Solutions, London, UK) with measurements obtained from the short-axis stack (figure 2A) using valve tracking on two corresponding long-axis cines (figures 2B-E). Papillary muscle and trabeculations were included in the mass measurement, using a semi-automated signal intensity-based thresholding technique. End-systolic and end-diastolic volumes were calculated thus: as appropriate, in systole or diastole, areas within manually traced endocardial borders (figure 2A) were calculated for each short axis cine, and multiplied by slice thickness to provide a slice volume. Slice volumes are measured along the ventricle from apex to the level of the mitral valve (figures 2B-E) and summed to calculate overall ventricular volume. Stroke volume and ejection fractions are calculated indirectly from end diastolic and end systolic volumes. Ventricular mass is quantified by measuring the area between endocardial and epicardial borders for sequential short axis cines (figure 2A), multiplying by the slice thickness and summing the volume of each slice. Total myocardial volume is multiplied by myocardial density (1.05g/ml) to provide a measure of myocardial mass. Volumes and mass were indexed to body surface area calculated using the Mosteller formula. Indexed volumetric data were: left ventricular (LV) mass; LV and right ventricular (RV) end-diastolic volumes; LV and RV end-systolic volumes; and LV and RV stroke volumes.

**Figure 1:**
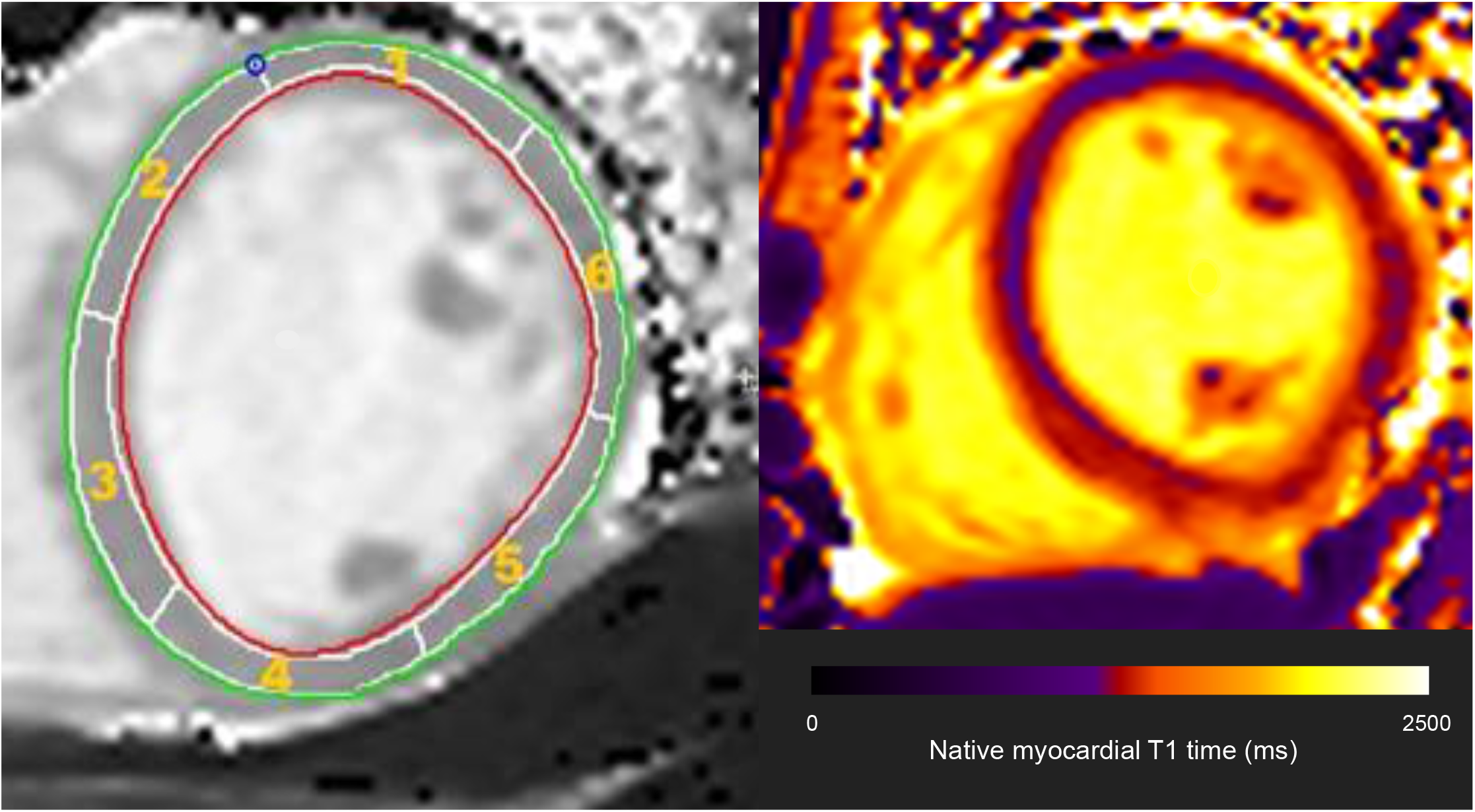
Showing the circumferential short axis segmentation of a mid-ventricular slice for purpose of native myocardial T1 time calculation, and representative T1 map. Red and green lines respectively represent the endocardial and epicardial borders. Each slice was divided into 6 segments with myocardial T1 time calculated for each segment. Segments with evidence of artefact were excluded from analysis, with the native myocardial T1 time for that participant then calculated as the mean T1 time of the remaining segments.

**Figure 2:**
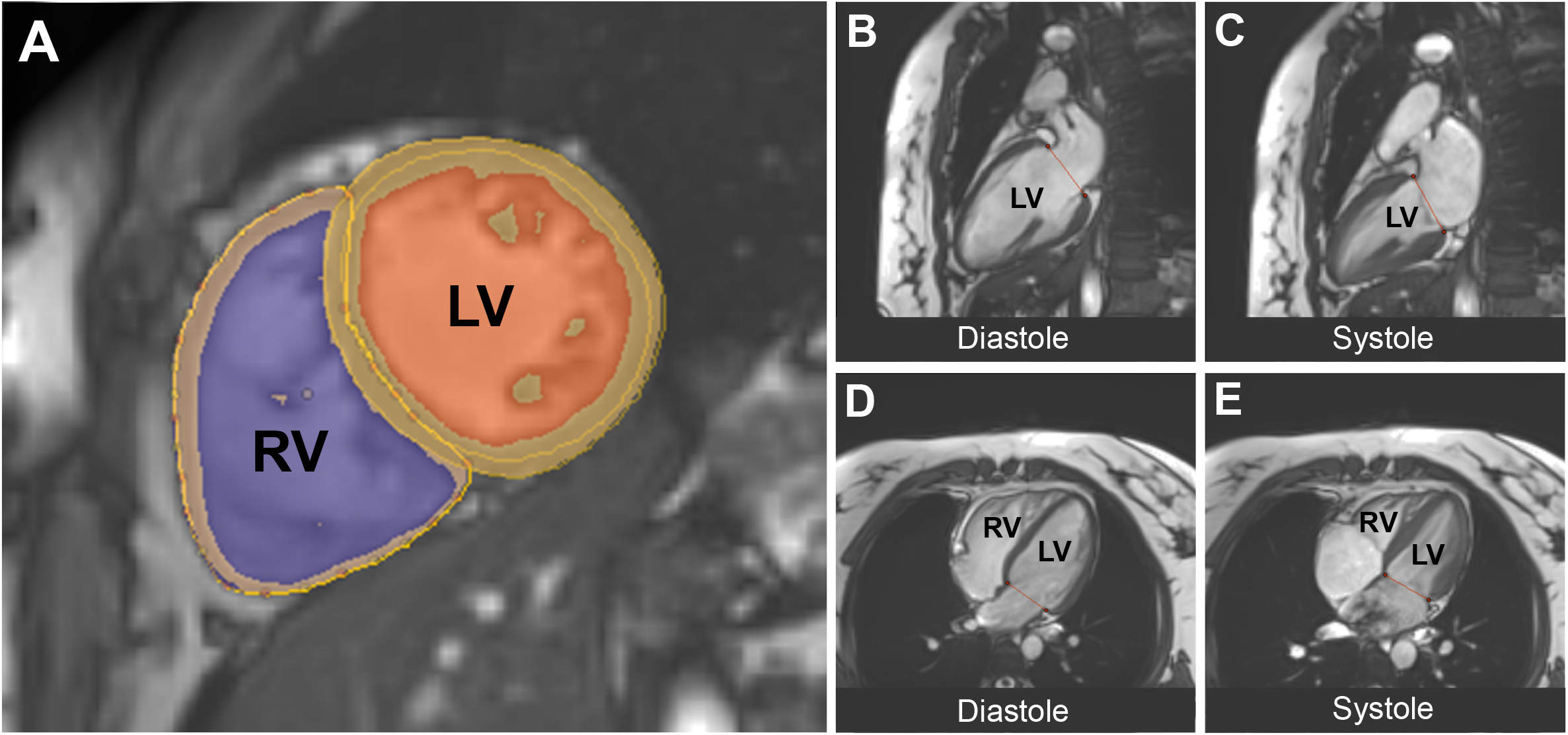
Showing the cardiac MR volumetric analysis. Figure 2A: Left ventricular short axis view with manually traced endocardial and epicardial borders. Semi-automated, signal intensity-based thresholding technique is used to include papillary muscles in the blood pool. **Figures 2B and 2C:** Mitral valve tracking in diastole and systole in the vertical long axis. **Figures 2D and 2E:** Mitral valve tracking in diastole and systole via a four-chamber view of the heart. Red line depicts level of mitral valve.

### Statistical analysis

The Shapiro-Wilk test of normality was used to assess continuous variables. Depending on the data being normally distributed, SCZ and HV groups were compared using either unpaired t-tests or Mann Whitney U tests, as appropriate. For nominal variables, Fisher’s exact test was used to compare groups. After recruitment was completed, we assessed for statistical matching of main factors recognized to be independently associated with cardiac dysfunction, including age, gender, ethnicity, smoking status, blood pressure, HbA1c, BMI and physical activity (see eAppendix 1 for more detailed description of physiological and environmental variables associated with cardiac dysfunction).

The primary outcome assessed was native myocardial T1 time. In addition, we conducted secondary analyses assessing cardiac structure (left ventricular mass) and function (left and right ventricular end-diastolic volumes, end-systolic volumes, stroke volumes, and ejection fractions). As these were exploratory, correction for multiple comparisons was not applied. Statistical analyses were performed on volumes and masses indexed to body surface area. Physiological variables were not included as covariates in statistical analysis of cardiac parameters owing to baseline statistical matching of these factors. Where we observed significant alterations in cardiac parameters between patients and controls, for normally distributed variables, the corresponding effect size (Cohen’s *d*) was calculated as the difference in mean cardiac MRI outcome between patients and controls divided by the pooled standard deviation. In line with international guidelines,^24^ we did not compare mean native T1 values with published normal ranges, as these outcomes are recognised to be scanner-specific.

Some patients were taking aripiprazole, which is a partial dopamine D2/3 agonist, in contrast to the other antipsychotics, which are all D2/3 antagonists.^25^ In case the inclusion of patients taking partial agonists biased the results, we conducted sensitivity analyses after excluding the subjects taking aripiprazole.

To further examine the putative link between antipsychotic use and myocardial fibrosis/inflammation, Spearman rank correlation coefficients were used to examine for an association between chlorpromazine equivalent dose/chlorpromazine dose years and native myocardial T1 time in patients. Moreover, because diabetes mellitus and hypertension are associated with development of myocardial fibrosis (eAppendix 1), we also set out to examine if these parameters were associated with alterations in native myocardial T1 time in patients. Spearman correlation coefficients were employed owing to the measure being robust to the influence of outliers. All statistical analysis was performed using SPSS software (version 25.0, Chicago, Illinois), for which statistical significance was defined as p < 0.05.

## Results

3 of the 17 SCZ participants dropped out of the study prior to scanning. Thus, 14 patients (mean age = 39.9 years; SD = 13.1 years) and 17 controls (mean age = 37.9 years; SD = 12.2 years) were included in final analyses. Further subject characteristics are described in table 1. SCZ and HV showed no significant differences in age, gender, smoking status, blood pressure, BMI, HbA1c, ethnicity, and physical activity measures (all p > 0.05). No participant met World Health Organisation criteria for type 2 diabetes mellitus or hypertension. For patients, duration of contact with mental health services ranged from 0.5-15 years. At the time of assessment, 3 patients were receiving aripiprazole, 3 olanzapine, and 8 clozapine (table 1 describes chlorpromazine equivalent doses and dose years). 6 of 31 participants (2 patients, 4 healthy volunteers) demonstrated evidence of artefact on 1 of the 6 segments making up the mid-ventricular slice for purpose of T1 mapping (figure 1). These segments were removed and native myocardial T1 calculated. Thus, 97% of ventricular segments were included in analyses. Continuous cardiac monitoring demonstrated that all participants were in sinus rhythm throughout the scans.

As shown in figure 3 and eTable 1, native myocardial T1 time was significantly longer in SCZ than HV with a large effect size (1212.38 ±21.23 ms vs 1190.96 ±26.01 ms, p = 0.02; Cohen’s *d* effect size = 0.89). Moreover, compared with HV, SCZ patients presented with lower indexed LV mass (d=0.76; p<0.05), smaller indexed LV end-diastolic volume (d=0.92; p=0.02), smaller indexed LV stroke volume (d=1.11; p=0.01), smaller indexed RV end-diastolic volume (p=0.02), and smaller indexed RV stroke volume (d=1.00; p=0.01). The difference was not significant in SCZ compared with HV for the following parameters: LV and RV ejection fraction, and indexed LV and RV end-systolic volume (all p > 0.05, eTable 2). Representative maps demonstrating differences between patients and controls for native myocardial T1 times are shown in figure 4.

**Figure 3:**
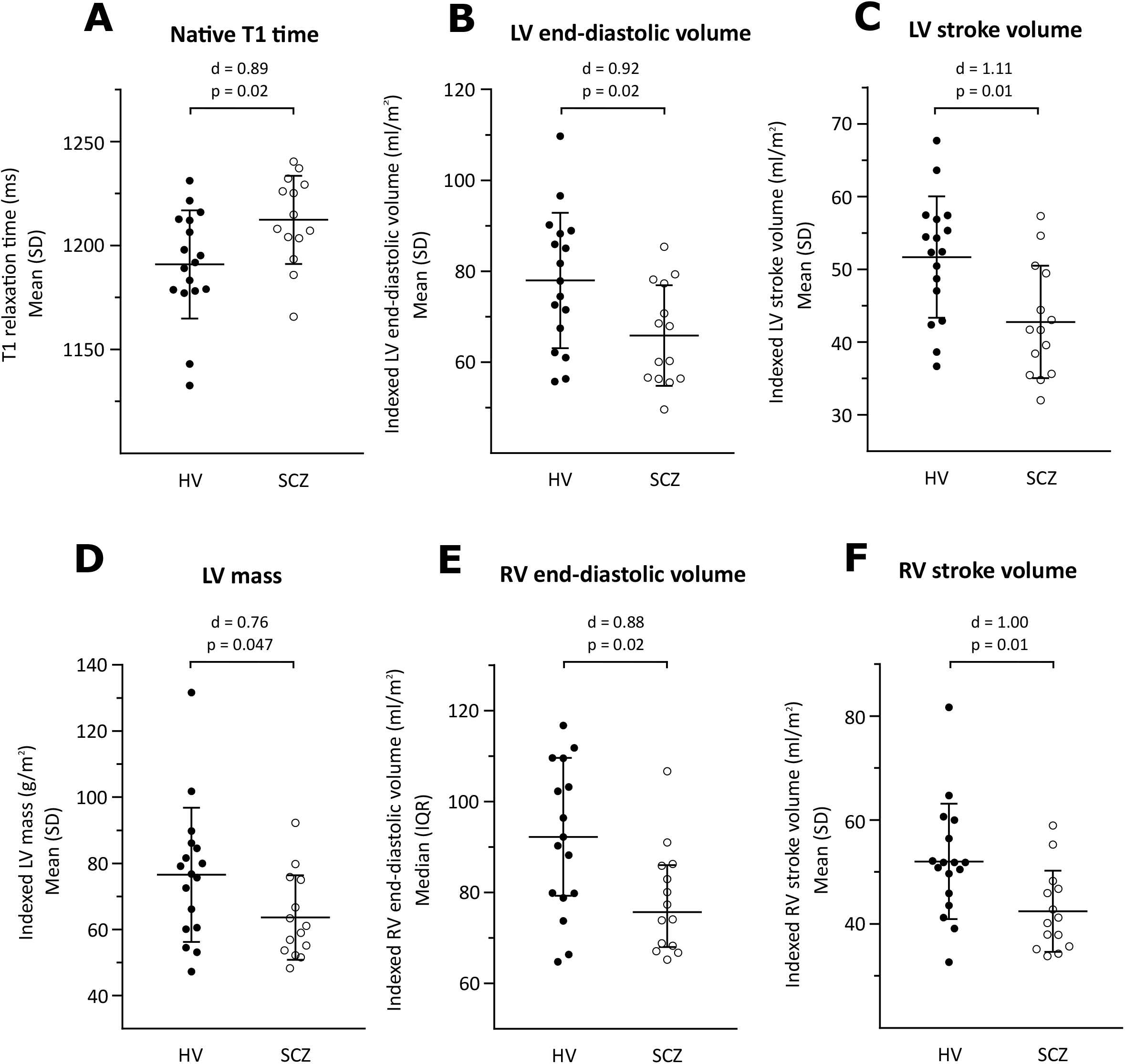
Cardiac structure and function in patients with schizophrenia (SCZ) and healthy volunteers (HV). **3A:** Native myocardial T1 time was longer in SCZ compared with HV (d=0.89; p=0.02); **3B:** Indexed left ventricular (LV) end-diastolic volume was smaller in SCZ compared with HV (d=0.92; p=0.02); **3C:** Indexed LV stroke volume was smaller in SCZ compared with HV (d=1.11; p=0.01); **3D:** Indexed LV mass was lower in SCZ compared with HV (d=0.76; p=0.047); **3E:** Indexed right ventricular (RV) end-diastolic volume was smaller in SCZ compared with HV (d = 0.88; p=0.02); **3F:** Indexed RV stroke volume was smaller in SCZ compared with HV (d=1.00; p=0.01). These findings are consistent with early diffuse myocardial fibrosis in patients with schizophrenia, d = Cohen’s d effect size for group mean difference. P values correspond to the unpaired t-test or Mann Whitney U test used to compare the 2 groups.

**Figure 4:**
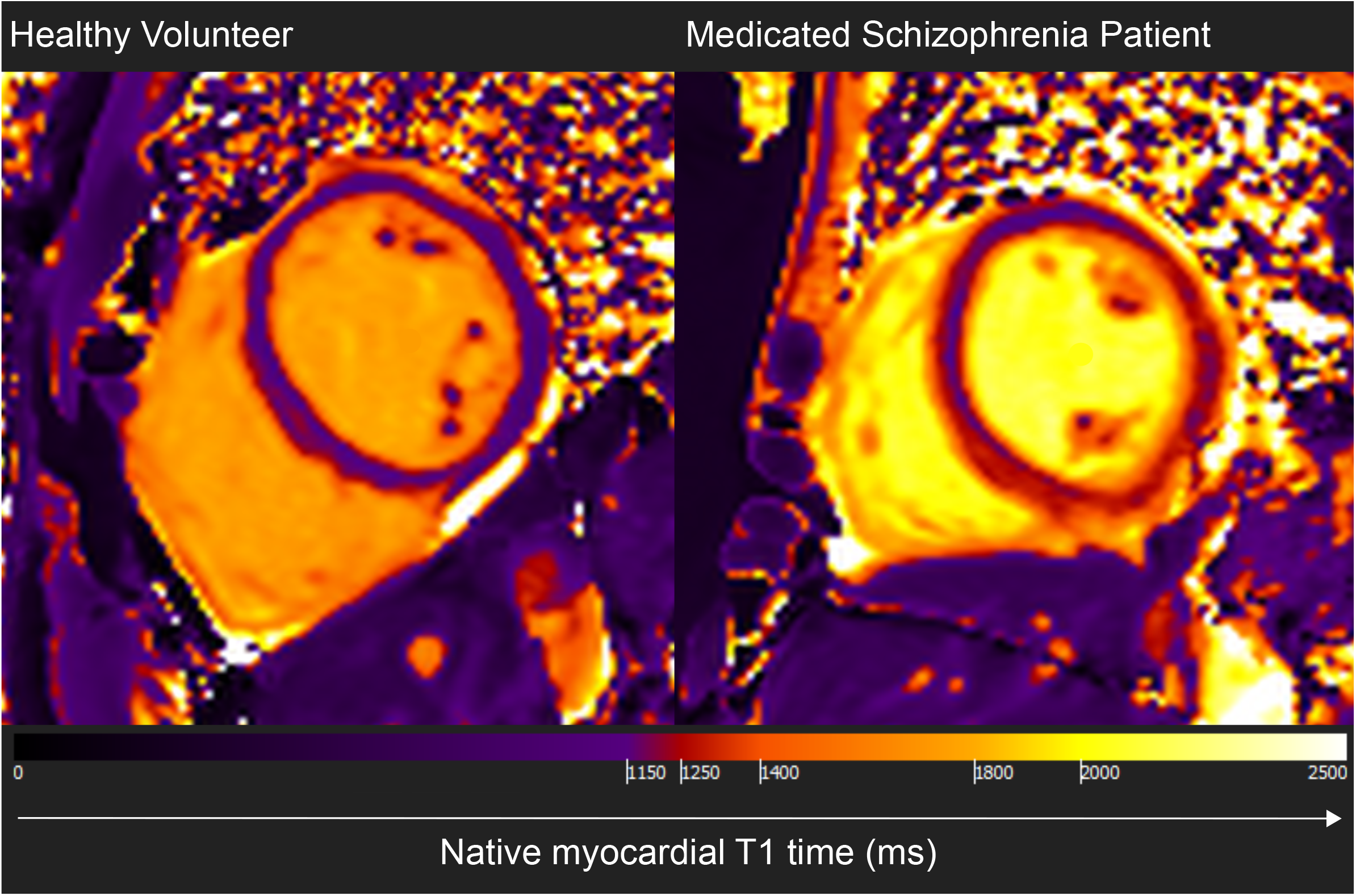
Representative cross-sectional mid-ventricular cardiac MRI images showing differences in native myocardial T1 maps for a healthy volunteer and a medicated patient with schizophrenia. Native myocardial T1 time is diffusely increased in schizophrenia compared with healthy volunteers.

Variation in T1 values between mid-myocardial segments and removal of 3% of ventricular segments (owing to identification of artefact) may have resulted in error in calculation of mean T1 time for some participants. However, re-inclusion of these 6 segments in the analysis did not significantly alter results, with T1 relaxation time remaining significantly longer in SCZ compared with HV (p = 0.02).

Removal of subjects taking aripiprazole (eTable 2) did not change results, except for indexed LV mass where the difference between patients and controls was no longer significant (p=0.12). No significant association was observed between T1 relaxation time and the following 4 variables: antipsychotic dose (Spearman r = 0.02; p = 0.94); duration of treatment (Spearman r = 0.10; p = 0.77), mean arterial pressure (Spearman r = 0.06; p = 0. 86), and HbA1c (Spearman r = −0.19; p = 0.63).

## Discussion

We observed that, compared with matched controls, native myocardial T1 was longer in antipsychotic-treated individuals with schizophrenia, who also exhibited lower indexed ventricular mass and smaller indexed ventricular and stroke volumes, with large effect sizes (d = 0.76-1.11). Subjects were matched for age, gender, ethnicity and BMI, and there were no statistical differences between the groups in terms of smoking status, blood pressure, HbA1c, and physical activity, indicating that our findings are unlikely to be explained by these cardiac risk factors.

The results of our study contrast with previous cardiac imaging studies in schizophrenia.^15,16,18^ Two of these studies,^15,16^ using transthoracic echocardiography to quantify cardiac structure and function, have demonstrated LV hypertrophy and reduced LV ejection fraction in patients, while we observed reductions in LV mass in patients, and no difference between groups with regards right/left ejection fraction. An explanation for these contrasting outcomes may relate to the sample characteristics of the patients included in analyses. Previous studies have either failed to match patients and healthy volunteers uniformly for demographics or cardiovascular risk factors,^16, 18^ or where matching for these parameters was achieved, confirmation of absence a diagnosis of diabetes mellitus in patients and controls was not confirmed with blood tests during the study.^15^ This is a limitation, owing to under-reporting of physical co-morbidities in schizophrenia.^26^ Moreover, these studies also failed to report the ethnicity of participants, which is of importance owing to the differential risk of metabolic disease and LV hypertrophy across ethnic groups (e.g. LV hypertrophy is more prevalent in individuals of black ethnicity).^27^ Echocardiographic evidence of cardiac disease in the context of metabolic syndrome and smoking (leading to ischaemic heart disease) is characterised by LV hypertrophy and potentially reduced ejection fraction.^28^ Thus, it is little surprise that these studies exhibited outcomes characteristic of a group with increased cardiovascular risk. However, in the current study, we matched patients and healthy volunteers for multiple cardiovascular risk factors and ethnicity, and in this context, absence of LV hypertrophy and EF reductions suggests that previously observed cardiac alterations in schizophrenia were likely the product of increased prevalence of cardiovascular risk factors in the patient group, alongside the added potential confound of different ethnic makeup of the two groups. The current study is also the first to observe reductions in end diastolic and stroke volumes in patients with schizophrenia compared with controls. Beyond the limitations of previous studies in matching patients with healthy volunteer groups, this could relate to differences in the method of assessing cardiac volumes, with MRI recognised as the more accurate measure compared with echocardiography.^19–22^

### Strengths and limitations

A strength of this study is the use of cardiac MRI to visualise the myocardium, which provides volumetric and functional cardiac measures that are not possible to derive from alternative cardiac imaging techniques, including assessments of myocardial tissue behaviour.^19^ Moreover, CMR is the gold standard for LV function and mass quantification,^20^ and has been demonstrated to provide better test-retest reliability than echocardiography,^21^ as well as lower inter- and intra-operator variability for a range of key measures.^22^ Thus, this study is the first to use the gold standard approach to measure cardiac function in schizophrenia research to date. A further strength is the matching of patients and controls for cardiac risk factors, thereby limiting the influence of confounders on outcome variables. Nevertheless, we cannot exclude residual confounding due to subtle sub-clinical metabolic or other variation.^29–31^ Moreover, it is recognised that measurement of certain variables (e.g. exercise) were based on self-reporting. The Copenhagen Physical Activity Score is not specifically designed for psychiatric populations, and individuals with schizophrenia can over-estimate the amount of physical activity in which they engage.^32^ This may have contributed to our finding of no significant difference in patients compared with controls with regards physical activity scores. However, a priori matching of groups in terms of BMI and exclusion of patients with cardiovascular comorbidity may by association have resulted in our observation of similar activity levels in the 2 groups. Future studies may consider including actigraphy to objectively quantify physical activity levels. Finally, although we measured BMI, we did not assess diet/abdominal obesity. However, metabolic alterations seen in schizophrenia or simply associated with poor diet, reduced exercise, and smoking would be expected to result in cardiac dysfunction typical of metabolic disease (such as LV hypertrophy^28^), the opposite to the pattern observed in this study. Indeed, although not statistically significant, in absolute terms BMI was raised in SCZ compared with HV (p = 0.31). However, increased BMI is associated with LV dilatation,^33^ while we observed the opposite in SCZ relative to HV. This suggests that residual confounding due to metabolic alterations or inaccurate reporting of lifestyle factors was not sufficient to alter outcomes, and indeed may have reduced the effect size magnitude of some parameters observed. Similarly, although patients and healthy volunteers were statistically matched for proportion of males and females (p = 0.50), there was still a relative increase in the proportion of females in the healthy volunteer group compared with patient group. Some,^34^ although not all,^35^ studies have observed prolonged native T1 in females compared with males. Moreover, smaller LV mass and ventricular volumes are observed in females compared with males.^34^ However, with a greater proportion of females in the HV group, one would expect this to result in relatively prolonged native T1 time and reductions in LV mass/volumes in HV, whereas we observed the opposite, suggesting this does not explain our findings. However, future studies with larger sample sizes that employ more assessments of cardiometabolic health (e.g. measurements of abdominal obesity) are indicated.

Removal of 3% of the mid-ventricular segments used to calculate mean T1 relaxation time (owing to identification of artefact) and variation in T1 values between the 6 mid-myocardial segments, may have altered the mean T1 calculation for some participants thereby influencing results. However, we are reassured that re-inclusion of these 6 segments in the analysis was not associated with any change in outcomes.

It should be recognised that native myocardial T1 time is not a specific marker of fibrosis. Future studies could employ further MRI measurements of myocardial fibrosis, such as post-contrast T1 (and extracellular volume calculation) mapping to provide additional validity, and T2 mapping to examine specifically for myocardial inflammation, although sensitivity for low-level oedema/inflammation is limiting.^36^

### Implications of our findings

Longer native myocardial T1 times in patients may reflect an early diffuse myocardial fibrosis and/or sub-clinical myocardial inflammation. Patients were taking a number of different antipsychotics including clozapine. Pre-clinical data suggest that both clozapine and non-clozapine antipsychotics can induce myocardial tissue alterations characteristic of fibrosis and inflammation, in a dose-dependent manner.^6–10^ Thus, our observations are compatible with antipsychotic treatment having a similar effect in humans, although future studies are required to examine if this is indeed a global effect of treatment, or specific to only certain drugs. When taken in the context of the broader outcomes of the current study, with longer native T1 times associated with lower ventricular mass, and smaller ventricular and stroke volumes, these findings are more compatible with a diffuse fibrotic process in patients resulting in reduced myocardial compliance, rather than a myocarditic process. Indeed, previous *in vivo* studies have observed that myocarditis is associated with increased LV mass,^37^ rather than our observation of a relative decrease in LV mass. Furthermore, although myocarditis is a recognised complication of clozapine, and 8/14 patients (57%) included in this sample were prescribed clozapine, patients included in the study were systemically well, with no history suggestive of myocarditis.

Our findings are in keeping with previous studies examining relationships between interstitial myocardial fibrosis and broader myocardial structure and function in the general population and in systemic inflammatory disease states such as rheumatoid arthiritis.^38^ These studies, after controlling for metabolic syndrome diagnoses, observe an inverse relationship between myocardial fibrosis and end-diastolic volumes and LV mass. In preclinical models of aging, the mechanism underlying reductions in LV mass in the context of fibrosis seem to relate to myocardial collagen fibre deposition and myocyte cell death occurring at a greater rate than concomitant hypertrophy in remaining myoctes.^39^ Reductions in end diastolic volumes are thought to be a consequence of the accumulation of collagen within the myocardium impacting its elasticity and limiting torsional recoil and ventricular suction.

Compared with the general population, people with schizophrenia are both twice as likely to have a diagnosis of CVD, and die as a consequence of CVD.^1^ Thus, the overall findings of this study, compatible with early diffuse myocardial fibrosis and/or sub-clinical myocardial inflammation in medicated patients with schizophrenia even in the absence of metabolic disease, could have significant clinical implications. For example, in various cardiac disease states, including non-ischaemic dilated cardiomyopathy and valvular pathology, myocardial fibrosis independently predicts both cardiovascular and all-cause mortality.^3^ If antipsychotic treatment in schizophrenia is indeed associated with a myocardial fibrotic process, then establishing the mechanism of these myocardial tissue alterations could potentially lead to identification of novel therapeutic approaches to reduce mortality rates in schizophrenia.

### Future directions

Taken in the context of previous preclinical studies that have observed myocardial fibrotic and inflammatory changes accompanying antipsychotic administration,^6–10^ the cardiac alterations observed in the current study may be the consequence of antipsychotic treatment. Further studies are therefore required to examine the potential link between antipsychotic medication and myocardial tissue alterations in humans by longitudinally assessing cardiac function in patients over time; when antipsychotic naïve and then during treatment.

However, the absence of significant correlations between antipsychotic dose/duration of treatment with native myocardial T1 time could be interpreted as suggesting that antipsychotics do not play a major role in the observed CMR alterations, although it is recognised that the study is not powered to examine this dose-response relationship. There is an emerging body of evidence suggesting that psychotic illness is independently associated with increased markers of oxidative stress and inflammation,^31,40^ systemic alterations that are associated with myocardial fibrosis and functional myocardial impairment.^14^ Examining myocardial alterations in antipsychotic naïve first episode psychosis would also provide insight into the role of physiological alterations that are putatively intrinsic to psychosis in altering myocardial structure and function.

Finally, studies examining the potential to attenuate accelerated myocardial fibrosis/inflammation in patients with schizophrenia may be required and could provide insight into novel therapeutic avenues with the potential to reduce mortality rates of patients with schizophrenia. Moreover, if antipsychotics are implicated in myocardial tissue alterations, examining antipsychotic dose-related cardiac alterations could guide clinicians with regards best practice in improving mental health without detrimentally impacting the physical health of patients. Furthermore, pharmacological interventions designed to target myocardial fibrosis/inflammation may be of therapeutic worth, and future studies exploring their use in antipsychotic treated individuals warranted. For example, drugs targeting the renin-angiotensin-aldosterone system have been observed to ameliorate myocardial fibrosis not only in preclinical models of hypertension^41^ and diabetes mellitus,^42^ but also in clozapine treated models.^43^

## Conclusions

Patients with schizophrenia taking antipsychotics show myocardial tissue alterations characteristic of an early diffuse fibrotic/inflammatory process that are not explained by other cardiac risk factors. These alterations could contribute to excess cardiac mortality rates observed in schizophrenia.

## Acknowledgments

We are grateful for the participation of the volunteers and to Professor Peter Jones, Cambridge, UK for his encouragement and helpful discussions. There was no financial compensation.

