## Supplementary Information for "Cardiac structure and function in patients with schizophrenia taking antipsychotic drugs: an MRI study"

| CMR System, pulse sequence, normal/reference ranges, changes in CMR-system related normal/reference ranges over time | |
| --- | --- |
| Clinical utility to assess for diffuse/global fibrosis | T1 (Native) mapping appropriate |
| T1, ECV and T2 mapping are typically performed at 1.5 or 3 Tesla (T). | 3 T scanner used |
| Normal/reference ranges: ‘a local reference range should be primarily used’ | N/A (comparing patients and controls, not with reference ranges) |
| Regularly repeated phantom-based quality control is recommended to ensure that status and stability of the CMR system have not changed significantly during the time between establishing normative values and clinical scanning. | Regular phantom-based quality control performed. No software update performed during study. |
| Imaging Protocols | |
| Native T1, T2, and T2* are measured in the absence of contrast agents | Achieved |
| Motivation and detailed instructions of patients are important to avoid incomplete breath-holds or motion artefacts. | Performed |
| Volume-selective B0 shimming focused on the heart is highly recommended at 1.5 T, and essential at 3 T. B1 (radiofrequency) volume shimming is recommended at 3 T. | Performed |
| T1 mapping | |
| For Look-Locker-based techniques (e.g. MOLLI), correction for readout-induced deflection of T1 relaxation is required (Look-Locker correction). | Achieved |
| Visualization and analysis | |
| Reporting clinicians should learn how to review source images and quality control maps to ensure registration/significant artefacts not present | Achieved |
| Maps may be displayed in colour if the colour look up tables are set according to site-specific ranges of normal | Achieved |
| The image reader should be trained with the local standards and with the analysis software package of choice and be aware of and familiar with the appearance of artefacts | Achieved |

**eTable 1: Further details regarding CMR-acquisition protocol and compliance with European Association for Cardiovascular Imaging (EACVI) guidance.**^1^

|  | Patients | Healthy Volunteers | Statistic in total sample | Statistic without patients on aripiprazole |
| --- | --- | --- | --- | --- |
| LV end-diastolic volume (ml)  Mean ±SD | 65.89 ±10.32 | 77.97 ±14.89 | t = 2.52  df = 29  p = 0.02 | t = 2.56  df = 26  p = 0.02 |
| LV end-systolic volume (ml)  Mean ±SD | 23.33 ±5.41 | 26.22 ±8.16 | t = 1.13  df = 29  p = 0.27 | T = 0.87  df =26  p = 0.39 |
| LV stroke volume (ml)  Mean ±SD | 42.76 ±7.72 | 51.69 ±8.33 | t = 3.07  df = 29  p = 0.01 | t = 3.54  df = 26  p = 0.002 |
| LV mass (g)  Mean ±SD | 63.62 ±12.77 | 76.54 ±20.29 | t = 2.07  df = 29  p = 0.0479 | t = 1.62  df = 26  p = 0.12 |
| RV end-diastolic volume (ml)  Median (IQR) | 75.69 (17.50) | 92.21 (30.28) | MWU  p = 0.02 | MWU  P = 0.03 |
| RV end-systolic volume (ml)  Median (IQR) | 35.51 (18.03) | 42.26 (8.53) | MWU  p = 0.06 | MWU  P = 0.13 |
| RV stroke volume (ml)  Mean ±SD | 42.47 ±7.78 | 52.06 ±11.11 | t = 2.72  df = 29  p = 0.01 | t = 2.80  df = 26  p = 0.01 |
| LV ejection fraction (%)  Mean ±SD | 64.86 ±4.88 | 66.94 ±5.27 | t = 1.13  df = 29  p = 0.27 | t = 1.71  df = 26  p = 0.10 |
| RV ejection fraction (%)  Mean ±SD | 54.41 ±4.72 | 54.43 ±4.69 | t = 0.01  df = 29  p = 0.99 | t = 0.77  df = 26  p = 0.45 |
| Relaxation time (ms)  Mean ±SD | 1212.38 ±21.23 | 1190.96 ±26.01 | t = 2.47  df = 29  p = 0.02 | t = 2.37  df = 26  p = 0.03 |

### **eTable 2:** **CMR-derived volumetry and functional assessments in the total sample (n=31) and with sensitivity analyses following removal of patients receiving aripiprazole (n=28).** The results remained essentially the same after excluding patients taking aripiprazole. Continuous data are presented as mean ±standard deviation (SD) where normally distributed, and as median (interquartile range (IQR)) where not. MWU: Mann Whitney U; RV: right ventricle; LV: left ventricle. Volumes/mass are indexed to body surface area.

**eAppendix 1**

**Supplementary Methods**

- SCZ and HV were assessed for statistical matching for main factors that are recognized to be independently associated with cardiac dysfunction, including age,^2^ gender,^2^ ethnicity,^3^ smoking status,^4^ blood pressure,^5^ HbA1c,^6^ BMI^5^ and physical activity^7^
- World Health Organisation criteria were used to define type 2 diabetes mellitus and hypertension.^8, 9^
- Brachial blood pressure measurements were performed following 5 minutes’ rest in accordance with European Society of Hypertension guidelines^20^
- Physical activity grading was based on the Copenhagen City Heart Study Leisure Time Physical Activity Questionnaire.^21^
- Chlorpromazine equivalent doses and dose-years were calculated as described by Andreason and colleagues.^22^
- Because diabetes mellitus^22^ and hypertension^23^ are associated with development of myocardial fibrosis, we set out to examine if these parameters were associated with alterations in native myocardial T1 time in patients.

**eAppendix 2**

### **Supplementary cardiac MRI Protocol**

- Myocardial tissue alterations were assessed in line with consensus guidelines (eAppendix 2)^1^ by measuring the longitudinal relaxation time constant of the myocardium (native myocardial T1 time) using a Modified Look-Locker Inversion recovery (MOLLI) sequence, 5s(3s)3s variant^10^ in a mid-ventricular short axis slice during breath-hold in end-expiration.^11^
- Each mid-ventricular slice was divided into 6 segments as per the American Heart Association model.^12^
- An epicardial and endocardial erosion offset of 10% was applied to the contours to ensure only myocardium was included.^11^
- A standard clinical protocol for assessing biventricular function and volumes was followed according to published international guidelines.^13^
- Slice volumes are measured along the ventricle from apex to the level of the mitral valve and summed to calculate overall ventricular volume.^14^
- Stroke volume and ejection fractions are calculated indirectly from end diastolic and end systolic volumes.^15^ Ventricular mass is quantified by measuring the area between endocardial and epicardial borders for sequential short axis cines, multiplying by the slice thickness and summing the volume of each slice.^14^
- Total myocardial volume is multiplied by myocardial density (1.05g/ml)^16^ to provide a measure of myocardial mass.
- Volumes and mass were indexed to body surface area calculated using the Mosteller formula.^17^

8.	World Health Organisation (WHO)/International Society of Hypertension (ISH) statement on management of hypertension. *World Health Organisation, International Society of Hypertension Writing Group* 2003.

9.	Definition and Diagnosis of Diabetes Mellitus and Intermediate Hyperglycaemia. *Report of World Health Organisation/International Diabetes Federation Consultation* 2006.
